# Bone marrow-derived CCR5-expressing macrophages promote venous thrombus resolution by up-regulating plasminogen activators and VEGF

**DOI:** 10.1101/2023.09.18.558360

**Authors:** Mizuho Nosaka, Yuko Ishida, Yumi Kuninaka, Akiko Ishigami, Akihiko Kimura, Akira Taruya, Atsushi Tanaka, Naofumi Mukaida, Toshikazu Kondo

**Author notes:** Corresponding to Toshikazu Kondo, MD & PhD, Department of Forensic Medicine, Wakayama Medical University, 811-1 Kimiidera, Wakayama 641-8509, Japan. Both authors equally contributed to this work (Mizuho Nosaka, Yuko Ishida).

## Abstract

**Background:** Deep vein thrombosis (DVT) is a common peripheral vascular disease arising from endothelial damage and can frequently result in pulmonary embolism. C-C chemokine receptor 5 (CCR5) is essentially involved in skin wound healing and pulmonary fibrosis, but the role of CCR5 still remains elusive.

**Methods:** Deep vein thrombus was induced by the ligation of the inferior vena cava (IVC) in wild-type (WT) mice and *Ccr5*^-/-^ mice, and thrombi were collected over time and various analyzes were performed.

**Results:** After ligation of the IVC in WT mice, a venous thrombus developed progressively until 5 days, and then resolved. Concomitantly, IVC ligation enhanced intrathrombotic gene and protein expression of *Ccr5* and its ligand, *Ccl5*, and both were expressed mainly by intrathrombotic macrophages. The same treatment of *Ccr5*^-/-^ mice resulted in significantly greater thrombus mass than WT mice. Moreover, the administration of a specific CCR5 inhibitor to WT mice recapitulated similar phenotypes as *Ccr5*^-/-^ mice while that of CCL5 caused the opposite phenotypes. In *Ccr5*^-/-^ mice, the production of intrathrombotic vascular endothelial growth factor (VEGF), tissue plasminogen activator (tPA), and urokinase-type plasminogen activator (uPA) from macrophages was significantly reduced and intrathrombotic recanalization was also suppressed compared to WT mice. Moreover, CCL5 enhanced *Vegf, Plat,* and *Plau* gene expression in WT**-**derived peritoneal macrophages by activating the ERK MAPK signaling pathway in vitro.

**Conclusions:** CCR5-deficient mice exhibited reduced expression of *Vegf, Plat,* and *Plau,* with concomitant attenuated neovascularization and reduced thrombus resolution. the Thus, the augmentation of the CCL5-CCR5 axis may be effective for DVT treatment by enhancing thrombolysis.

## Introduction

Venous thromboembolism (VTE) is the most common vascular cause of death in the world after coronary heart disease and ischemic stroke (1) and its morbidity and mortality are comparable to those of Alzheimer’s disease and diabetes (2). VTE encompasses deep vein thrombosis (DVT) and its two major sequelae, pulmonary thromboembolism (PE) and post-thrombotic syndrome. Dislodgment of thrombus in pulmonary vein causes PE while post-thrombotic syndrome develops when thrombus formation and resolution damage vein walls, thereby causing venous hypertension (3, 4).

Blood flow turbulence, endothelial injury, and hypercoagulability are main causes of DVT (5). In contrast, several lines of evidence additionally indicate the involvement of inflammatory responses in DVT resolution (6–9). Venous thrombus resolves through a process of organization and recanalization, the process that is similarly observed in granulation tissue formation in wound healing. Our previous observations on the roles of proinflammatory cytokines in wound healing process (10, 11, 12) prompted us to explore the roles of various cytokines in DVT resolution. Indeed, interferon (IFN)-γ, could play a detrimental role in thrombus resolution by suppressing metalloproteinase (MMP)-9 and vascular endothelial growth factor (VEGF) expression (13) while tumor necrosis factor (TNF)-α and interleukin (IL)-6 could promote thrombus resolution (14, 15). These observations suggest that DVT resolution is governed by the complicated network of various proinflammatory cytokines.

Inflammatory cell recruitment is an important component of thrombus resolution as well as granulation tissue formation in skin wound healing. The absence of CXCR2, a receptor for potent neutrophilic chemokines, impaired thrombus resolution in mice (16), together with decreased intrathrombotic neutrophil and monocyte recruitment, and reduced neovascularization (17). Moreover, evidence is accumulating to indicate the essential involvement of a monocyte-trophic C-C chemokine, CCL2, in thrombus resolution (18–20). However, it still remains elusive on the roles of other chemokines/chemokine receptors in DVT formation and/or resolution.

CCR5 is a seven-transmembrane G protein-coupled receptor and primarily expressed in monocytes, macrophages and T cells (21, 22), although its expression is also observed in cells of different tissues (23, 24). It is a promiscuous receptor that binds with three different ligands such as CCL3, CCL4, and CCL5, and induces chemotactic cell migration upon binding with its ligands. It plays fundamental roles in the inflammatory response by recruiting cells to sites of inflamed sites (25). Moreover, accumulating evidence demonstrated that CCR5 was involved in tissue repair or fibrosis by inducing migration of bone-marrow-derived cells including endothelial progenitor cells (EPCs) (11) and fibrocytes (26). Furthermore, CCR5 engagement also results in the activation of several signal transduction cascades such as protein kinase B (PKB or Akt), NF-κB and ERK pathways (27). Indeed, the CCL3-CCR5 axis has a protective role in inflammation-induced aneurysm formation by activating p38 MAPK pathway (28). However, it remains to be clarified on the pathophysiological roles of CCR5 in DVT.

Here, we investigated the pathophysiological roles of CCR5 in DVT by using *Ccr5^-/-^* mice. We revealed that *Ccr5^-/-^*mice exhibited retarded resolution of the venous thrombosis together with reduced expression of plasminogen activators (PAs) and VEGF. Moreover, the CCL5-CCR5 axis enhanced the gene expression of PAs and VEGF in macrophages by activating ERK. Thus, the CCL5-CCR5 axis may be able to promote venous thrombus resolution.

## Materials and Methods

### Reagents and antibodies (Abs)

Recombinant mouse CCL3, CCL4 and CCL5 were purchased from R&D Systems (Minneapolis, MN). Maraviroc (a specific CCR5 inhibitor, PZ0002) was obtained from Sigma-Aldrich (Saint Lois, MO). The following monoclonal Abs (mAbs) and polyclonal Abs (pAbs) were used for immunohistochemical and double-color immunofluorescence analyses: rat anti-mouse F4/80 mAb (Dainippon Pharmaceutical Company, Osaka, Japan), rat anti-mouse CD31 mAb (BD Biosciences, San Jose, CA), goat anti-mouse CCR5 mAb, rabbit anti-tPA pAbs, rabbit anti-uPA pAbs, goat anti-mouse MMP-2 pAbs, goat anti-mouse MMP-9 pAbs, anti-mouse VEGF pAbs, goat anti-mouse CCL5 pAbs (Santa Cruz Biotechnology, Dallas, TX), rabbit anti-myeloperoxidase (MPO) pAbs (Neomarkers, Fremont, Fremont, CA), cyanine dye 3-conjugated donkey anti-rabbit IgG or anti-goat IgG pAbs, FITC-conjugated donkey anti-rat IgG pAbs (Jackson Immunoresearch Laboratories, West Grove, PA). Western blot analysis was performed by using the following Abs: rabbit anti-mouse p38 MAPK pAbs, rabbit anti-mouse JNK pAbs, rabbit anti-mouse ERK mAb, rabbit anti-mouse phosphorylated (p)-p38 MAPK mAbs, rabbit anti-mouse p-JNK pAbs, rabbit anti-mouse p-ERK mAbs, rabbit anti-mouse GAPDH pAbs (Cell Signaling Technology, Danvers, MA).

### Mice

As wild-type (WT) mice, pathogen-free 8- to 10-wk-old male C57BL/6 mice were obtained from Japan SLC (Shizuoka, Japan) in this study. Age- and sex-matched *Ccr5*^−/−^ mice were backcrossed with C57BL/6 mice for more than 8 generations and were used in these experiments (28). All mice were housed individually in cages under the specific pathogen-free conditions during the experiments. All animal experiments were approved by the Committee on Animal Care and Use in Wakayama Medical University (No. 1152).

### IVC ligation-induced deep vein thrombus model

Intravenous thrombus formation was induced as described previously (15). In brief, after deep anesthesia with intraperitoneal injection of triple agents (medetomidine hydrochloride, midazolam, and butorphanol), a 2-cm incision was made along the abdominal midline followed by IVC ligation with 3-0 silk suture. In some experiments, WT mice were intraperitoneally given a CCR5 inhibitor (Maraviroc, 10 mg/kg in 200 μl PBS) at 1 day before IVC ligation and at 1, 3 and 6 days after the ligation. At the indicated time intervals after IVC ligation, mice were euthanized by an overdose of isoflurane, to harvest intravenous thrombi for various analyses including weight measurement.

### Generation of bone marrow (BM) chimeric mice

The following BM chimeric mice were prepared: male *Ccr5^-/-^* BM to female WT mice, male WT BM to female *Ccr5^-/-^* mice, male WT BM to female WT mice, and male *Ccr5^-/-^* BM to *Ccr5^-/-^* mice (11). BM cells collected from the femurs of donor mice by aspiration and flushing. Recipient mice were irradiated at 12 Gy and then intravenously received 5 x10^6^ BM cells from donor mice in a volume of 200 μl sterile PBS under anesthesia. Thereafter, the mice were housed in sterilized microisolator cages and were fed normal chow and autoclaved hyperchlorinated water for 6 weeks. To verify successful engraftment and reconstitution of the BM in the transplanted mice, genomic DNA was isolated from peripheral blood and tail tissues of each chimeric mouse 6 weeks after BM transplantation with NucleoSpin® Tissue Kit (MACHEREY-NAGEL, Duren, Germany). Then, we performed PCR to detect the *Sry* gene contained in the Y chromosome (Forward primer, 5’-TTGCCTCAACAAAA-3’; Reverse primer, 5’-AAACTGCTGCTTCTGCTGGT-3’).

The amplified PCR products were fractionated on a 2% agarose gel and visualized by ethidium bromide staining. After durable BM engraftment was confirmed, intravenous thrombus formation was induced in the recipient mice as described above.

### Histopathological analyses

At the indicated time intervals after IVC ligation, thrombi were harvested, fixed in 4% formaldehyde buffered with PBS (pH 7.2), and paraffin-embedded sections (4 μm thick) were made. The sections were stained with hematoxylin and eosin or Masson trichrome solution.

### Immunohistochemical analyses

Immunostaining was automatically performed by Ventana Discovery® XT (Ventana Medical Systems, Inc., AZ). The primary Abs were diluted with the blocking buffer (PBS containing 1% normal serum corresponding to the secondary Abs and 2% bovine serum albumin to reduce nonspecific reactions. Thereafter, the sections were reacted with rabbit anti-MPO pAbs, rat anti-F4/80 mAb (20 μg/ml), goat anti-CCR5 pAbs (2 μg/ml), rat anti-CD31 (5 μg/ml), goat anti-mouse MMP-2 pAbs (4 μg/ml) or goat anti-mouse MMP-9 pAbs (4 μg/ml) at 37°C for 60 min. After incubation with biotinylated anti-rabbit IgG pAbs (1.3 μg/ml), anti-rat IgG pAbs (0.65 μg/ml) or anti-goat IgG pAbs (1.3 μg/ml) at 37°C for 60 min, the DAB Map kit (Ventana Medical Systems, Inc., AZ) was used to visualize the antigens for all stains. Subsequently, all slides were counterstained with hematoxylin and lithium carbonate to visualize the nuclei (15).

### Determination of intrathrombotic neovessel density and MMP-positive cells

Intrathrombotic vascular density and MMP-positive cells were semi-quantitatively evaluated as previously described (14). Briefly, intrathrombotic CD31^+^ areas with tube-like formation were counted in 5 high-power fields (×1,000) as vascular density, as previously described (14). MMP-2^+^ cells or MMP-9^+^ cells in five high power fields (×1,000) were counted within the thrombus, and the total numbers in the five fields were combined. All measurements were performed by an examiner without prior knowledge of the experimental procedures.

### A double-color immunofluorescence analysis

In order to determine the cell type expressing CCR5, CCL5, or VEGF, double-color immunofluorescence analyses were performed by Ventana Discovery XT (15). Briefly, deparaffinized sections were incubated with PBS containing 1% normal donkey serum and 1% BSA to reduce nonspecific reactions, as described previously (29). Thereafter, the sections were further incubated in the combination of anti-F4/80 mAb (20 μg/ml) and anti-CCR5 or anti-CCL5 pAbs (2 μg/ml), or anti-VEGF pAbs (1 μg/ml) and anti-CCR5 pAbs. After incubation with Cy3-conjugated and FITC-conjugated secondary pAbs (each at a concentration of 15 μg/ml) at 37°C for 60 min, nuclei were stained using 4’,6-diamidino-2-phenylindole (DAPI, Roche Diagnostics) according to the manufacturer’s instructions.

### IVC blood flow measurement by a laser Doppler

Microvascular IVC blood flow was evaluated by laser Doppler imaging (OMEGAFLO FLO-C1 BV, OMEGAWAVE) as described previously (13). Blood flow through the exposed IVC region of the interest was assessed, at three time points; immediately after laparotomy, immediately after the ligation, and at the harvest. The intensities were reported as the percentage of the baseline blood flow of each animal, in order to ensure consistency.

### Real-time RT-PCR

Real-time RT-PCR was performed as described previously (28). Briefly, total RNAs were extracted from tissue samples (100 μg) using ISOGEN (Nippon Gene, Toyama, Japan) according to the manufacturer’s instructions, and 5 μg of total RNAs were reverse-transcribed into cDNA at 42°C for 1 h in 20 μl reaction mixture containing mouse Moloney leukemia virus reverse transcriptase (PrimeScript, TAKARA BIO, Shiga, Japan) with random 6 primers (TAKARA BIO). Thereafter, generated cDNA was subjected to real-time PCR analysis using SYBR^®^ *Premix Ex Taq*™ II kit (TAKARA BIO) with the sets of specific primers (Table 1). Relative quantity of the target gene expression to β-actin gene was measured by comparative Ct method.

**Table 1.**
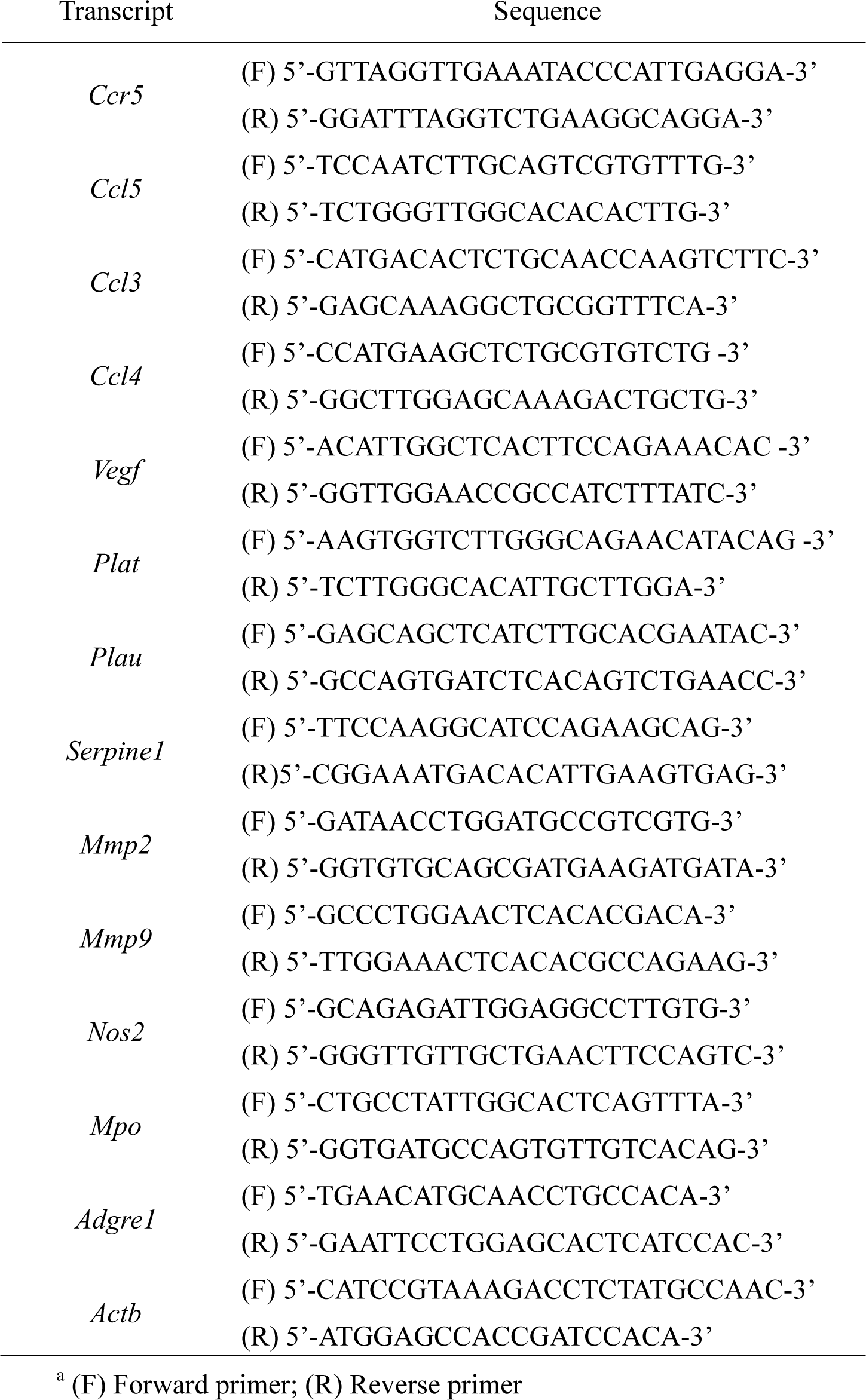
Sequences of the primers used for real-time RT-PCR.

### ELISA

At the indicated time intervals, thrombus samples were obtained and homogenized with 0.3 ml of PBS (pH 7.2) containing Complete Protease Inhibitor Mixture (Roche Diagnostics). The homogenates were centrifuged at 12,000 g for 15 min. The supernatants were subjected to ELISA by using commercially available kits for VEGF, (VEGF, Mouse, ELISA Kit, Quantikine M, MMV00, R&D Systems, Minneapolis, MN), tPA (Mouse tPA ELISA Kit, PA92, Oxford Biomedical Research, Rochester Hills, MI), uPA (Active mouse uPA Functional Assay Kit, PL92, Oxford Biomedical Research, Rochester Hills, MI), PAI-1 (Moue PAI-1 ELISA Kit, ELM-PAII, RayBiotech, Peachtree Corners, GA) or CCL5 (Mouse CCL5/RANTES ELISA Kit, Quantikine ELISA, MMR00, R&D Systems, Minneapolis, MN) according to the manufacturer’s instructions. The detection limits for each method were follows: VEGF, > 3.0 pg/mL; tPA, > 0.1 ng/mL; uPA, > 0.125 ng/mL; PAI-1, > 80 pg/mL; CCL5, > 2 pg/mL. Total protein in the supernatant was measured with a commercial kit (BCA Protein Assay Kit; Pierce, Tokyo, Japan) using BSA as a standard. The data were expressed as VEGF (pg/ml)/total protein (mg/ml) for each sample.

### Cell culture

WT mice were i.p. injected with 2 ml of 3% thioglycollate (Sigma-Aldrich), to obtain intraperitoneal macrophages (macrophage purity > 95%) 3 days later as described previously (28). The resultant cells were judged to consist of more than 95% macrophages as determined by a flow cytometer using anti-F4/80 Ab and were suspended in antibiotic-free DMEM medium containing 10% FBS and incubated at 37°C in three 6-well cell culture plates. Two hours later, nonadherent cells were removed, and the medium was replaced. After the cells were incubated for 24 h in the presence of recombinant mouse CCL5 (20 ng/ml) together with or without an ERK inhibitor, PD098059 (4 μg/ml), the cells were applied to subsequent analyses.

### Western blotting

The obtained macrophages were homogenized with a lysis buffer (20 mM Tris-HCl (pH 7.6), 150 mM NaCl, 1% Triton X-100, 1 mM EDTA) containing Complete protease inhibitor cocktail (Roche Diagnostics) and were centrifuged to obtain lysates. The lysates (equivalent to 30 µg protein) were electrophoresed in a 10% SDS-polyacrylamide gel and were transferred onto a nylon membrane. After the membrane was sequentially reacted with optimally-diluted primary Abs and HRP-conjugated secondary Abs, the immune complexes were visualized using ECL system (Amersham Biosciences). The band intensities were measured using NIH Image Analysis Software version 1.50a (National Institutes of Health).

### Measurement of prothrombin time (PT) and activated partial thromboplastin time (APTT)

Blood samples were taken with 3.8% citrate solution and centrifuged to obtain plasma samples. PT and APTT of citrated plasma samples were measured by using COAGSEARCH (A&T, Ashburn, VA) according to the manufacturer’s instructions.

### Statistical analysis

Data were expressed as the mean ± SEM. The normality and equal variance of all data could be confirmed by the use of Klomogorov–Smirnov test and Bartlett one, followed by subsequent analyses. For the comparison between WT and Ccr5^−/−^ mice at multiple time points, 2-way ANOVA followed by Dunnett’s *post hoc* test was used. To compare the values between the 2 groups, a two-sided unpaired Student’s *t* test was performed. In the experiment to examine the effects of CCL5 stimulation with several inhibitors on peritoneal macrophages, 1-way ANOVA followed by Dunnett’s *post hoc* test was used. *P* < 0.05 was considered statistically significant. All statistical analyses were performed using Statcel3 software.

### Ethics

All animal experiments were approved by the Committee on Animal Care and Use at Wakayama Medical University (No. 1152). All methods were performed in accordance with relevant guidelines and regulations including the ARRIVE guideline.

## Results

### Intrathrombotic *Ccr5* and *Ccl5* expression after inferior vena cava (IVC) ligation

The expression of *Ccr5* mRNA was detectable in the thrombus at 1 day after IVC ligation, increasing progressively as thrombi aged (Fig. 1A). Consistently, intrathrombotic CCR5^+^ cells were immunohitochemically detected in the thrombus (Fig. 1B), and a double-color immunofluorescence analyses demonstrated that CCR5 were mainly expressed on F4/80^+^ macrophages (Fig. 1C). We further noticed the gene expression of CCR5 ligands (CCL3, CCL4 and CCL5) in thrombi later than 1 day after IVC ligation (Fig. 1, D-F), but the gene expression levels of *Ccl3* and *Ccl4* was not enhanced until 14 days after IVC ligation (Fig. 1, D and E). On the contrary, the expression of *Ccl5* mRNA was markedly augmented and reached a maximum at 5 days after IVC ligation (Fig. 1F). Consistently, intrathrombotic CCL5 protein contents were increased later than 5 days after IVC ligation (Fig. 1G) and CCL5 protein was expressed mainly in intrathrombotic F4/80^+^ macrophages (Fig. 1H). These observations would imply the involvement of the CCL5-CCR5 axis in the formation and/or resolution of deep vein thrombi in an autocrine or paracrine manner. In order to explore the pathophysiological roles of CCR5 in IVC ligation-induced venous thrombus formation, we compared thrombus formation between WT and *Ccr5*^−/−^ mice. In WT mice, venous thrombi developed progressively until 5 days, resolving gradually thereafter (Fig. 1, I and J). Later than 5 days after IVC ligation, thrombi masses were larger in *Ccr5*^−/−^ mice than in WT ones (Fig. 1, I and J). Intrathrombotic collagen area was further remarkably enhanced in *Ccr5*^−/−^ mice, compared with WT ones (Fig. 1, K and L). We next determined IVC blood flow as the degree of intrathrombotic recanalization, which was presumed to be essential for thrombus resolution (13–15). Actually, IVC blood flow was significantly lower at 10 and 14 days in *Ccr5*^−/−^ mice than in WT mice (Fig. 1M). Consistently, intrathrombotic neovascularization was depressed in the absence of CCR5 as evidenced by reduced CD31^+^ areas in *Ccr5*^−/−^ mice 14 and 21 days after IVC ligation, compared with WT ones (Fig. 1, N and O). Moreover, treatment with a CCR5 antagonist, Maraviroc, increased thrombus mass (Fig. 1P) and reciprocally depressed blood flow (Fig. 1Q) 6 days after IVC ligation, compared with vehicle treatment. On the contrary, recombinant CCL5 (rCCL5) administration reduced significantly the thrombus weights and accelerated blood flow recovery (Fig. 1, R and S) when it was started 1 day after IVC ligation. These observations demonstrated that the CCL5-CCR5 axis could promote the resolution of venous thrombi.

**Figure 1.**
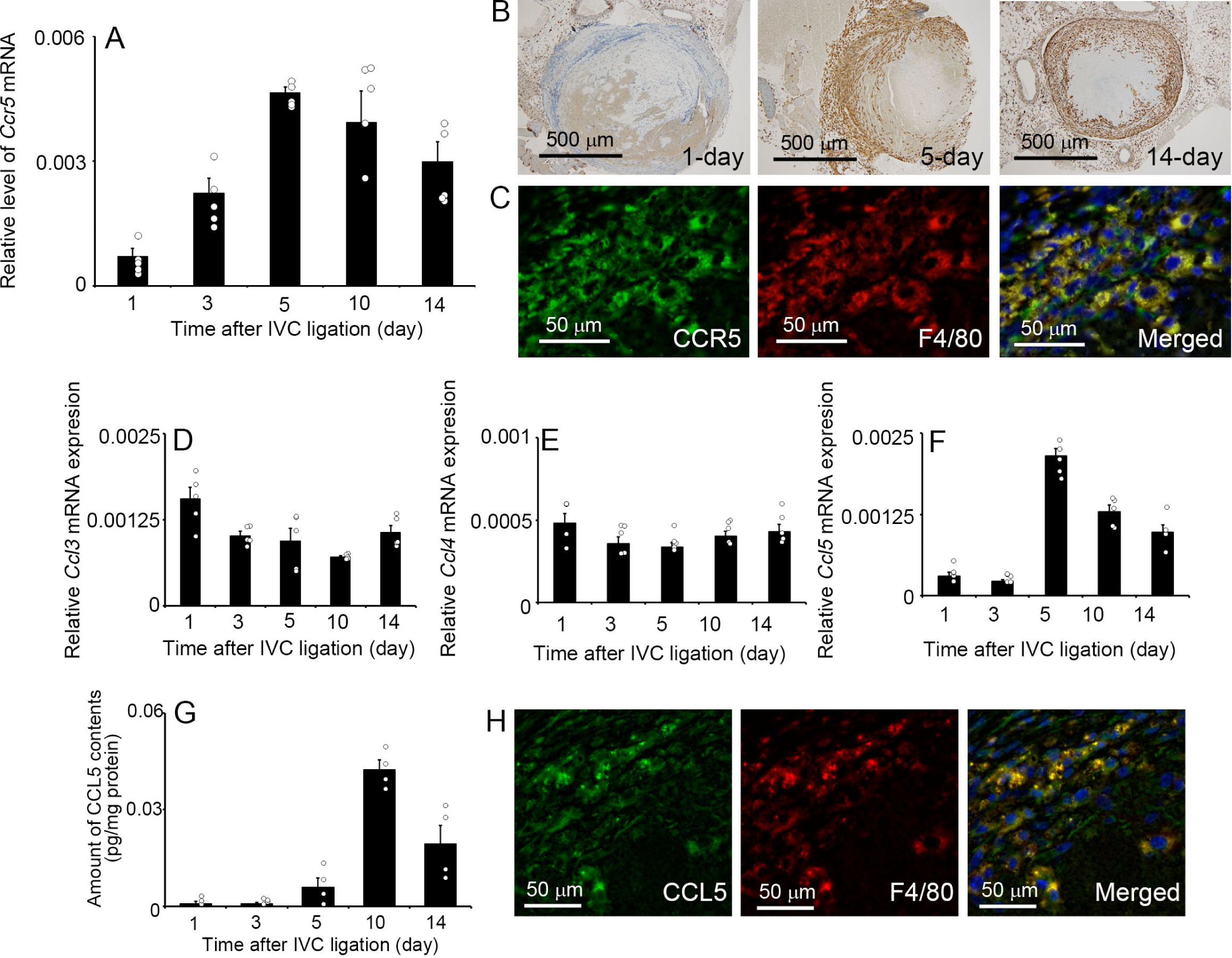
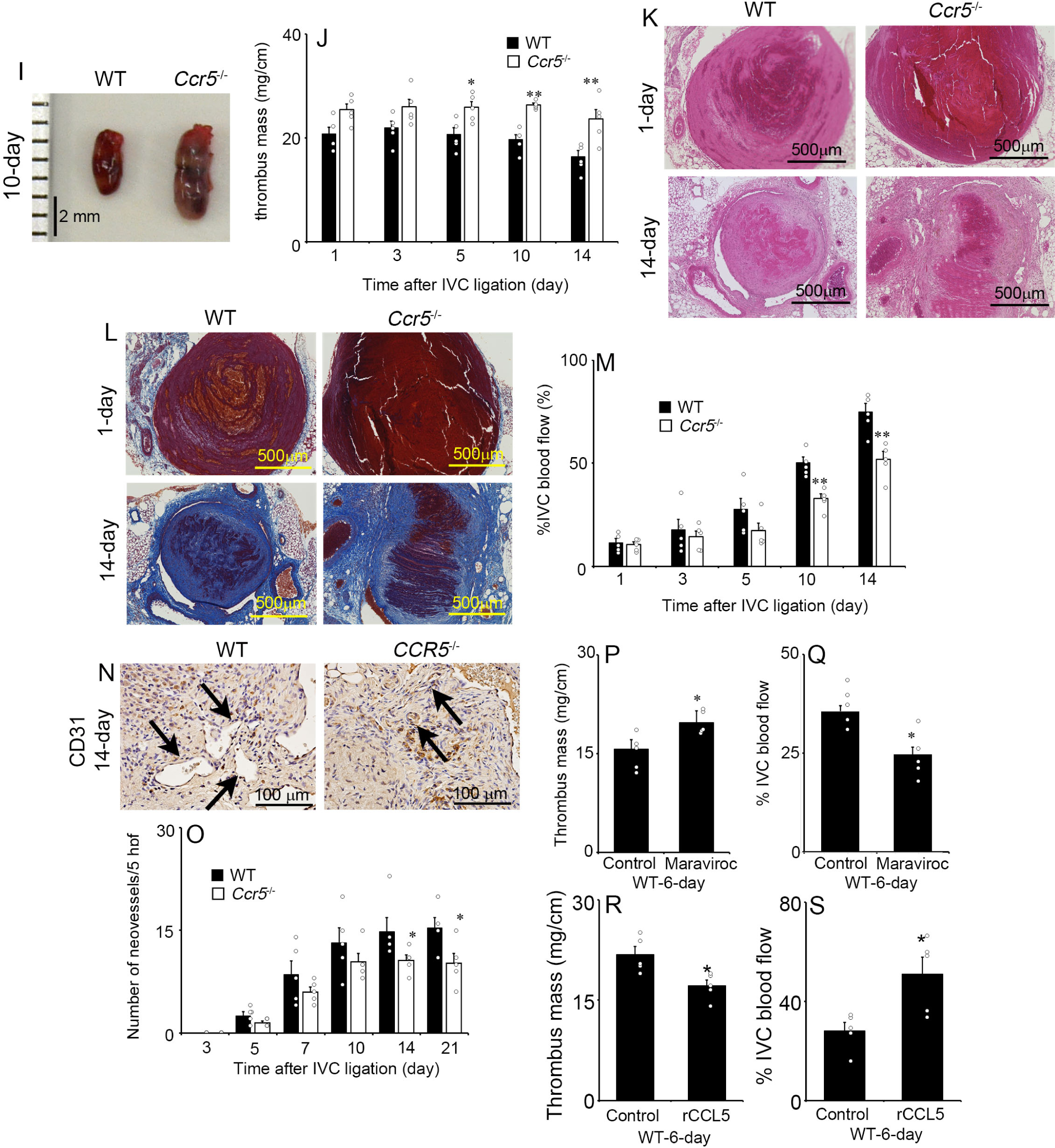
Intrathrombotic expression of CCR5 and CCL5 in WT mice after IVC ligation. (A) *Ccr5* gene expression was determined by real-time RT-PCR. All values represent mean ± SEM (n = 5). (B) Immunohistochemical analysis was performed using anti-CCR5 in thrombus samples from WT mice. Representative results from 6 independent experiments are shown. (C) A double-color immunofluorescence analysis of CCR5-expressing cells in the thrombus at day 5. The samples were immunostained with the combination of anti-CCR5 and anti-F4/80. (D-F) The gene expression of *Ccl3* (D), *Ccl4* (E) and *Ccl5* (F) were examined by real-time RT-PCR. All values represent mean ± SEM (n = 5). (G) Intrathrombotic contents of CCL5 in WT mice after IVC ligation. All values represent mean ± SEM (n = 4). (H) A double-color immunofluorescence analysis of CCL5-expressing cells in the thrombus at day 5. The samples were immunostained with the combination of anti-CCL5 and anti-F4/80. (I-S) IVC ligation-induced deep vein thrombus in WT and *Ccr5*^-/-^ mice. (I) Macroscopic appearance of thrombi in WT and *Ccr5*^-/-^ mice 10 days after IVC ligation. Representative results from 6 independent animals are shown. (J) Thrombus mass of WT and *Ccr5*^-/-^ mice at the indicated time intervals after IVC ligation. All values represent the mean ± SEM (n = 5). \*\**P* < 0.01; \**P* < 0.05, vs. WT. (K and L) Histopathologic analysis of thrombi obtained from WT and *Ccr5*^-/-^ mice. Thrombi were stained with HE (K) or Masson trichrome solution (L). (M) The measurement of IVC blood flow by laser Doppler imaging. All values represent the mean ± SEM (n = 5). \*\**P* < 0.01, vs. WT. (N) Immunohistochemical analysis was performed using anti-CD31 on thrombus samples from WT and *Ccr5*^-/-^ mice. Representative results from 6 independent experiments are shown (day 14). Arrowheads indicate the CD31^+^ areas with tube-like formation. (O) CD31^+^ vascular areas were determined (n = 5). \**P* < 0.05, vs. WT. (P) Thrombus mass of vehicle- and Maraviroc-treated WT mice at day 6 after IVC ligation. Values represent the mean ± SEM (n = 5). \**P* < 0.05, vs. vehicle-treated controls. (Q) The measurement of IVC blood flow by laser Doppler imaging. All values represent the mean ± SEM (n = 5). \**P* < 0.05, vs. vehicle-treated controls. (R) Thrombus mass of PBS- and rCCL5-treated WT mice at day 6 after IVC ligation. Values represent the mean ± SEM (n = 5). \**P* < 0.05, vs. PBS-treated controls. (S) The measurement of IVC blood flow by laser Doppler imaging. All values represent the mean ± SEM (n = 5). \**P*<0.05, vs. PBS-treated controls.

### The absence of CCR5 had no effects on intrathrombotic leukocyte recruitment

Coagulation and subsequent leukocyte infiltration are observed after IVC ligation similarly as observed on other types of tissue injury and can contribute to thrombus formation (30–32). However, *Ccr5*^−/−^ mice exhibited similar coagulation functions as WT mice as evidenced by the absence of prolonged PT and APTT (Fig. S1). Immunohistochemical analyses revealed that intrathrombotic MPO^+^ neutrophil and F4/80^+^ macrophage numbers after IVC ligation were similar between *Ccr5*^−/−^ and WT mice (Fig.2, A and C). Supporting this notion, *Ccr5*^−/−^ and WT mice exhibited similar levels of gene expression of *Mpo* (a neutrophil marker) and *Adgre1* (a macrophage marker) during the whole course of thrombus formation after IVC ligation (Fig. 2, B and D). Together, the absence of CCR5 might have few, if any, effects on intrathrombotic leukocyte recruitment.

**Figure 2.**
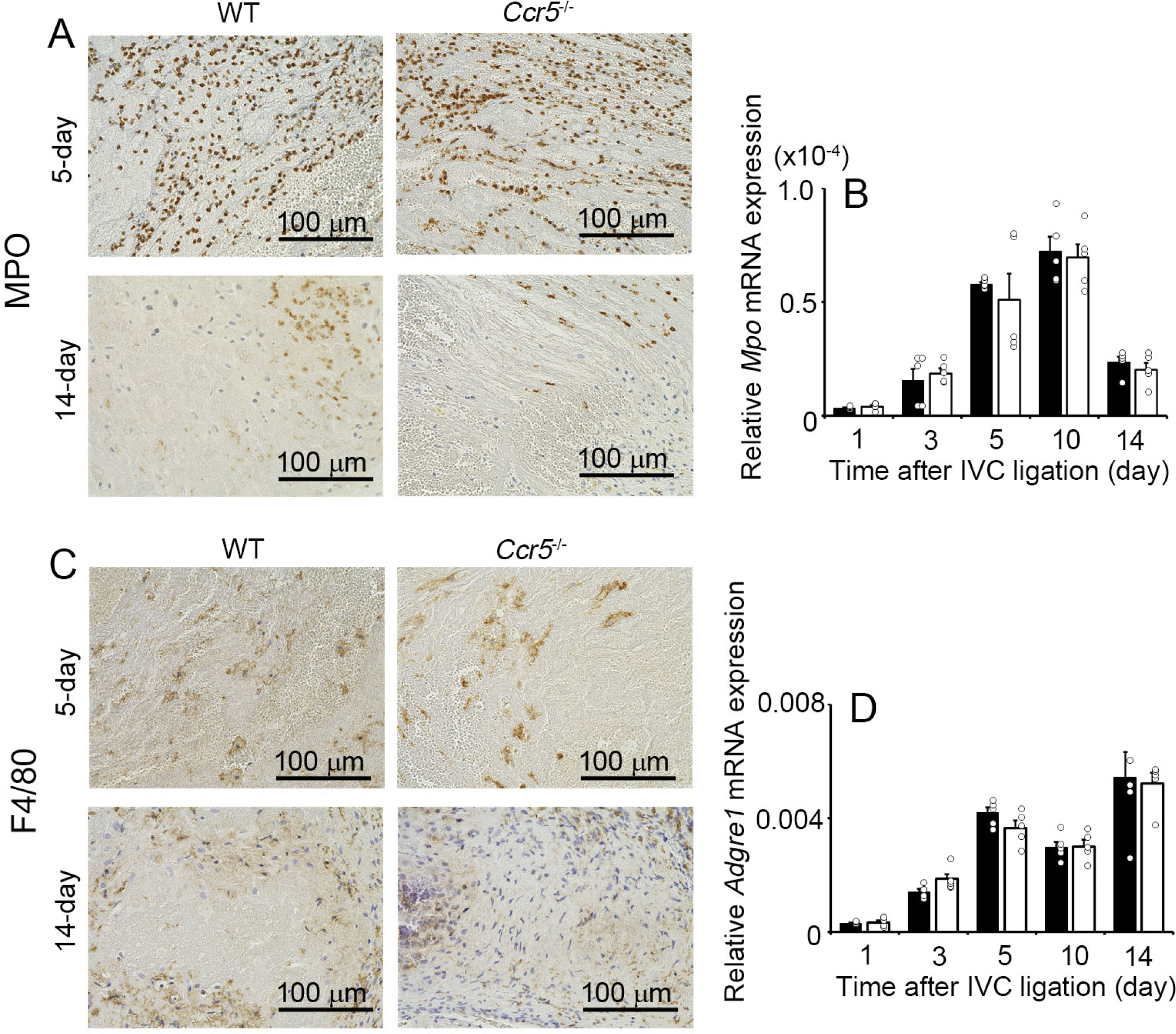
Enumeration of intrathrombotic leukocytes in WT and *Ccr5*^-/-^ mice. (A and C) Immunohistochemical analysis was performed using anti-MPO (A) or anti-F4/80 (C) in thrombus samples from WT and *Ccr5*^-/-^ mice. Representative results from 6 independent experiments are shown. The gene expression of *Mpo* (B) and *Adgre1* (D) were determined. All values represent the mean ± SEM (n = 5).

### Contribution of CCR5^+^ BM-derived cells to thrombus resolution

Evidence is accumulating to indicate that BM-derived cells could be detected in the thrombus after IVC ligation (33, 34), and that CCR5 is expressed by BM-derived and non-BM-derived cells (11, 35). Hence, we explored the contribution of BM-derived CCR5^+^ cells to thrombus resolution by using BM chimeric mice generated between WT and *Ccr5^-/-^* mice. Both WT and *Ccr5^-/-^* mice transplanted with *Ccr5^-/-^* mouse-derived BM cells exhibited impaired thrombus resolution as evidenced by larger thrombus mess and delayed blood flow recovery compared with the respective recipients of WT BM cells (Fig. 3, A to E). Consistently, neovascular areas were smaller in both WT and *Ccr5^-/-^*mice transplanted with *Ccr5*^-/-^ mouse-derived BM cells than in the respective WT BM recipients at 14 days after IVC ligation (Fig. 3, F to G). These observations would imply an essential contribution of BM-derived CCR5^+^ cells to thrombus resolution.

**Figure 3.**
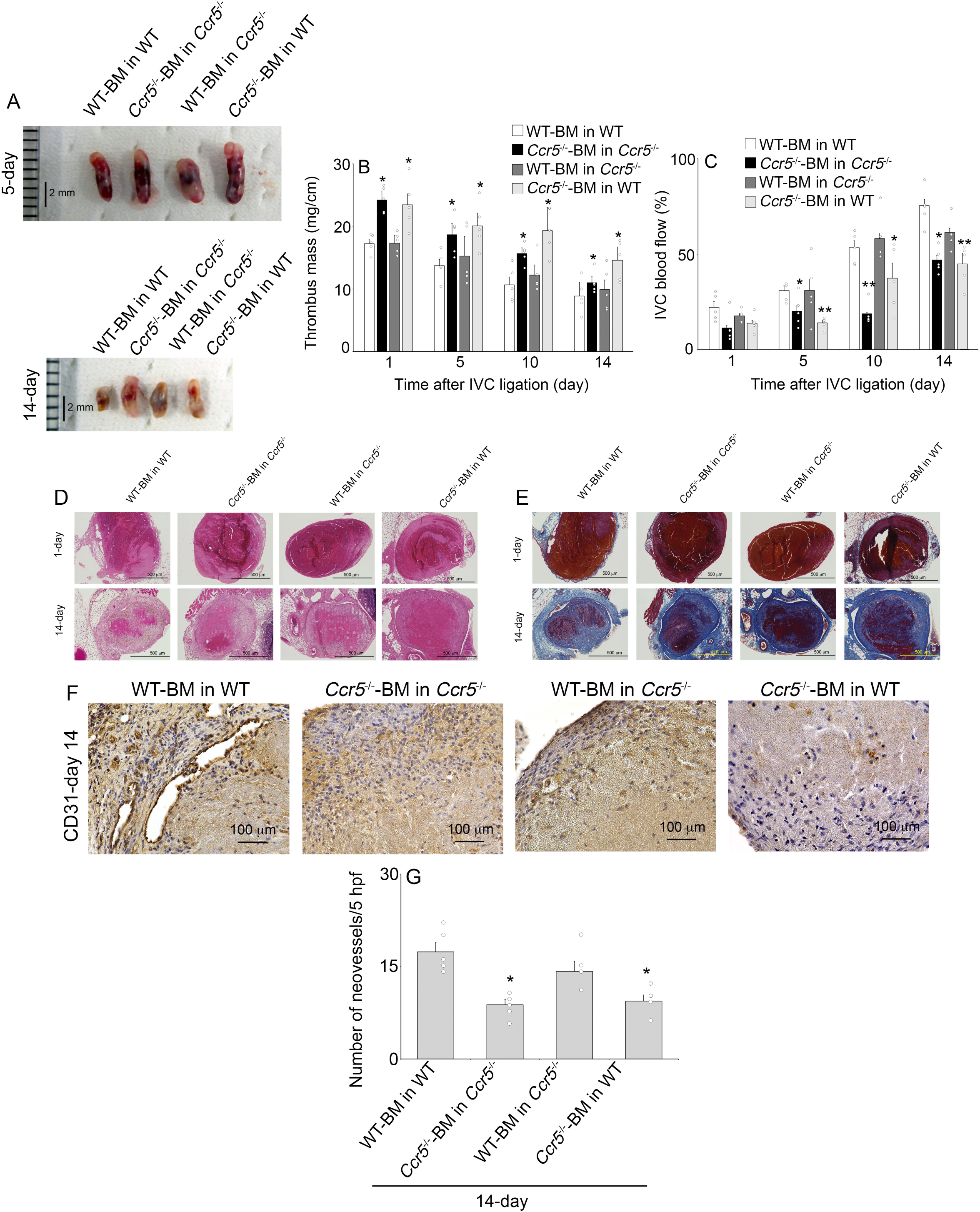
Effects of BM transplantation on thrombus formation of recipient mice transplanted with BM cells from WT and *Ccr5*^-/-^ donors. (A) Macroscopic appearance of thrombi in BM chimeric mice at day 5 and 14 after IVC ligation. Representative results from 6 independent animals are shown. (B) Thrombus mass of WT and *Ccr5*^-/-^ mice transplanted with WT or *Ccr5*^-/-^ mouse-derived BM cells at the indicated time intervals after IVC ligation. All values represent the mean ± SEM (n = 5). \**P*<0.05, vs. WT recipient with WT-BM. (C) The measurement of IVC blood flow by laser Doppler imaging. All values represent the mean ± SEM (n = 5). \*\**P* < 0.01, vs. WT recipient with WT-BM. (D and E) Histopathologic analysis of thrombi obtained from BM chimeric mice. Thrombi were stained with HE (D) or Masson trichrome solution (E). (F) Immunohistochemical analysis was performed using anti-CD31 in thrombus samples. Representative results from 6 independent experiments are shown. (G) CD31^+^ vascular areas were determined. All values represent the mean ± SEM (n = 5). \**P* < 0.05, vs. WT recipient with WT-BM.

### Reduced intrathrombotic expression of VEGF, tissue-type PA (tPA), and urokinase-type PA (uPA) in *Ccr5^−/−^*mice

Given the indispensable role of VEGF in neovascularization, we next examined the gene expression of intrathrombotic *Vegf* in WT and *Ccr5*^-/-^ mice. Later than 5 days after IVC ligation, intrathrombotic *Vegf* gene expression was markedly enhanced in WT mice but the enhancement was depressed in *Ccr5*^-/-^ mice (Fig. 4A). Consistently, intrathrombotic VEGF protein levels were significantly lower in *Ccr5*^-/-^ mice than in WT mice (Fig. 4B). Moreover, a double immunofluorescence analysis demonstrated that VEGF proteins were detected in CCR5^+^ cells in thrombi (Fig. 4C) and that VEGF^+^ cells expressed a macrophage marker, F4/80 (Fig. S2). These observations would indicate that CCR5 deficiency could reduce VEGF expression by macrophages, eventually reducing intrathrombotic neovascularization. We next examined intrathrombotic gene expression of tPA (*Plat*), uPA (*Plau*), plasminogen activator inhibitor-1 (*Pai1*), and matrix matrix metalloproteinases (*Mmps*), the molecules that are presumed to have crucial roles in thrombus resolution (13–15). The intrathrombotic gene expression and protein contents of tPA and uPA but not those of PAI-1 were significantly attenuated in *Ccr5*^-/-^ mice, compared with WT ones (Fig. 4, D to I). In line with our previous study (15), we detected both tPA and uPA proteins in intrathrombotic F4/80^+^ macrophages and CCR5-expressing cells (Fig. 4, J to M). On the contrary, there were no significant differences in intrathrombotic gene and protein expressions of MMP-2 and MMP-9 between WT and *Ccr5*^-/-^ mice (Fig. 4, N to S). Together, CCR5^+^ macrophage-derived tPA and uPA might contribute to thrombus resolution.

**Figure 4.**
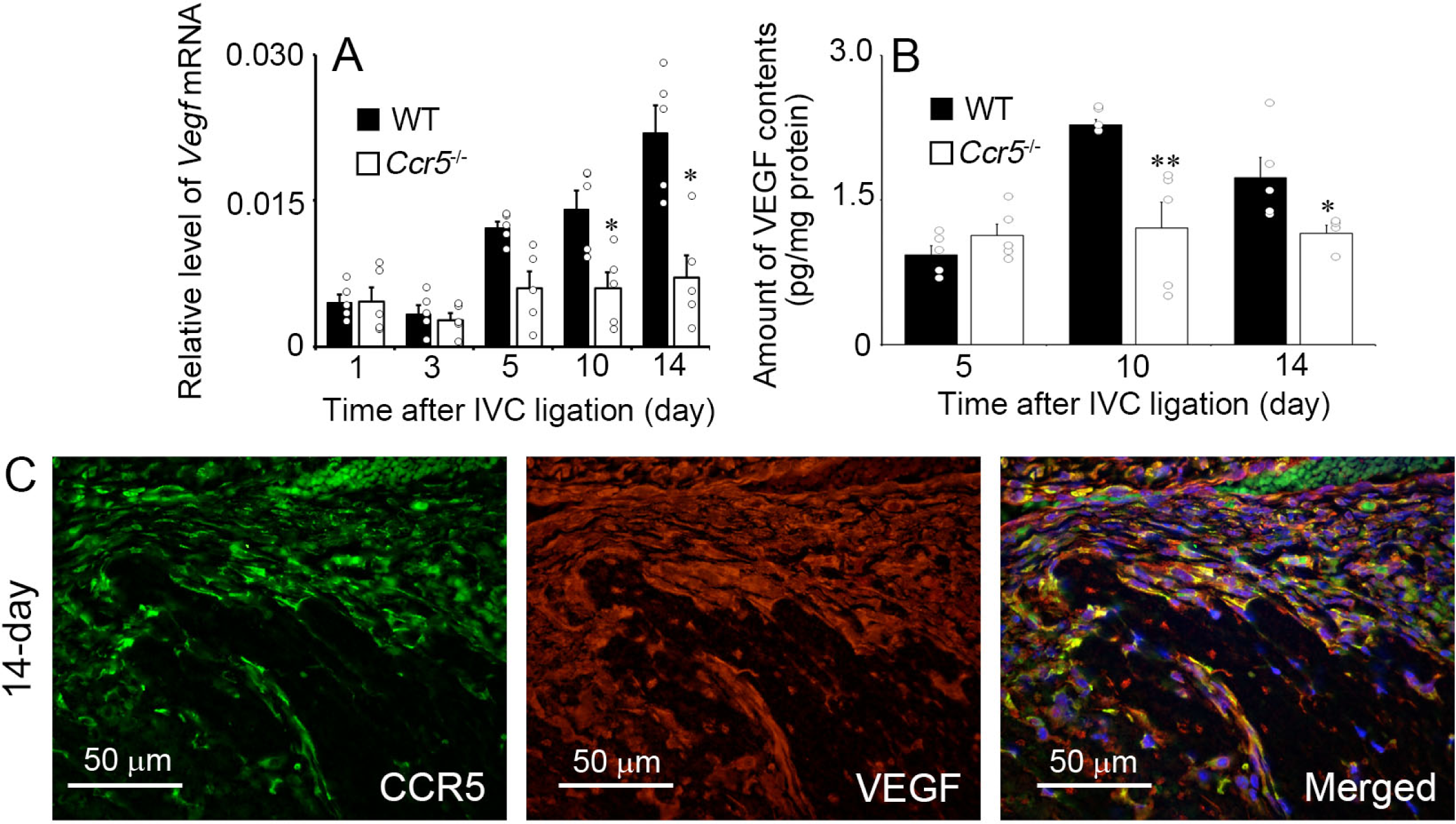
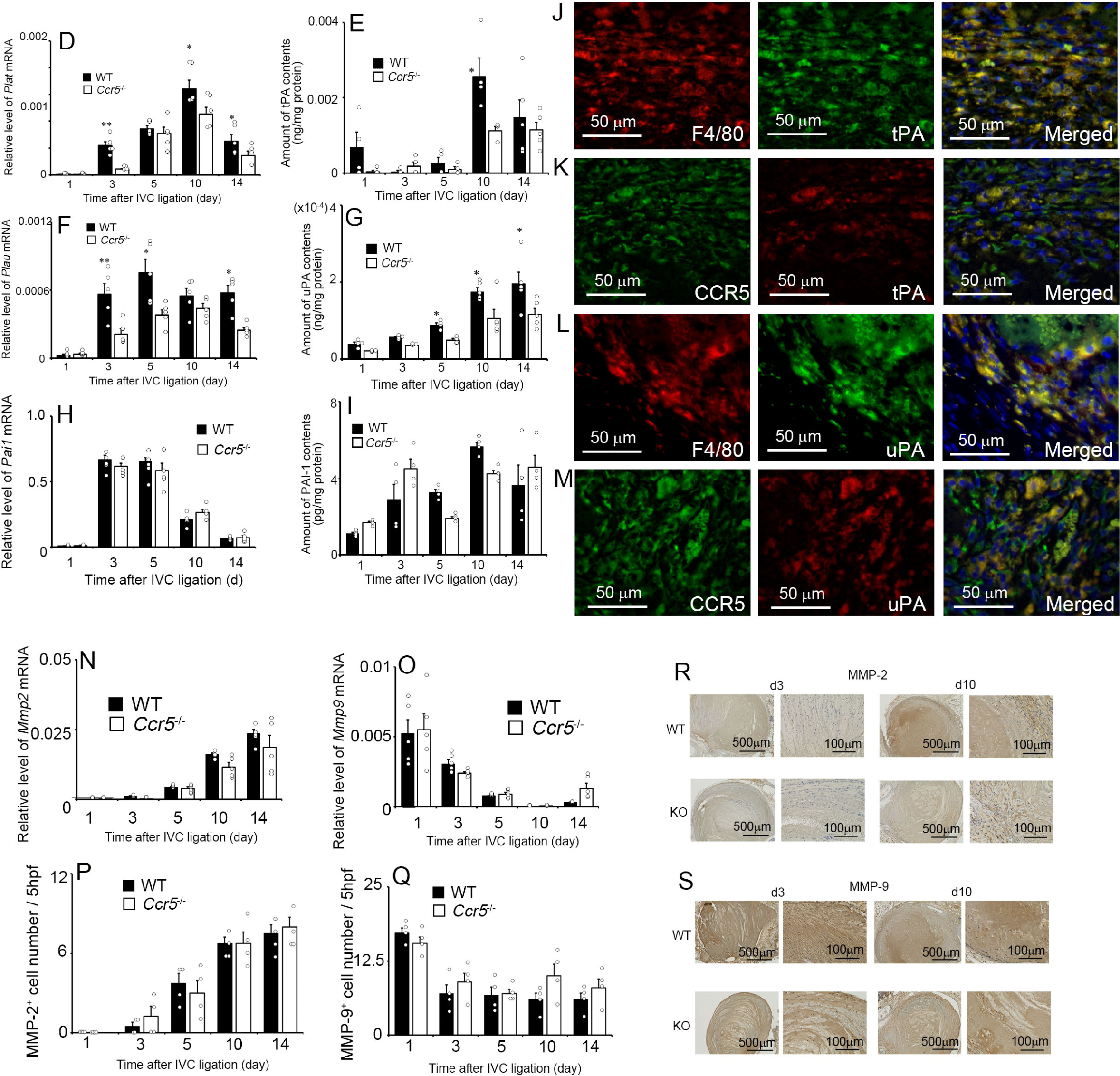
Evaluation of intrathrombotic recanalization. (A) Intrathrombitic gene expression of *Vegf* in WT and *Ccr5*^-/-^ mice. All values represent the mean ± SEM (n = 5). (B) Intrathrombotic VEGF protein contents were determined at the indicated time intervals after IVC ligation. All values represent the mean ± SEM (n = 5). \*\**P* < 0.01; \**P* < 0.05, vs. WT. (C) A double-color immunofluorescence analysis of VEGF-expressing cells in the thrombus at day 14. The samples were immunostained with the combination of anti-VEGF and anti-CCR5. (D-S) (D, F, and H) Intrathrombotic gene expression of *Plat* (D) and *Plau* (F), and *Pai1* (H) in WT and *Ccr5*^-/-^ mice. All values represent the mean ± SEM (n = 5). \*\**P* < 0.01; \**P* < 0.05, vs. WT. (E, G, and I) Intrathrombotic contents of tPA (E, n = 5), uPA (G, n = 5), and PAI-1 (I, n = 4) in WT and *Ccr5*^-/-^ mice. All values represent the mean ± SEM. \**P* < 0.05, vs. WT. (J-M) A double-color immunofluorescence analysis of tPA- and uPA-expressing cells in the thrombus at day 10. The samples were immunostained with the combination of anti-tPA and anti-F4/80 (J) or anti-CCR5 (K). The samples were also immunostained with the combination of anti-uPA and anti-F4/80 (L) or anti-CCR5 (M). (N and O) Intrathrombotic gene expression of *Mmp2* (N) and *Mmp9* (O) in WT and *Ccr5*^-/-^ mice. All values represent the mean ± SEM (n = 5). (P-S) Immunohistochemical analysis was performed using anti-MMP-2 or anti-MMP-9 in thrombus samples from WT and *Ccr5*^-/-^ mice. The numbers of MMP-2^+^ (P) or MMP-9^+^ cells (Q) were determined. All values represent the mean ± SEM (n = 4). Representative results from 6 independent experiments are shown in R (MMP-2) and S (MMP-9).

### ERK signal pathway regulated the gene expression of *Vegf*, *Plat* and *Plau* in macrophages

Our in vivo observations suggest that the CCL5-CCR5 axis might induce the gene expression of *Vegf*, *Plat* and *Plau* in intrathrombotic macrophages. To validate this hypothesis, we examined the effects of the CCL5-CCR5 axis on the gene expression of *Vegf*, *Plat* and *Plau* in peritoneal macrophages. CCL5 significantly enhanced the gene expression of *Vegf*, *Plat* and *Plau* in WT-derived but not *Ccr5*^-/-^-derived macrophages (Fig. 5, A-C). Moreover, CCL5 significantly enhanced the phosphorylation of ERK but not p38 MAPK and JNK (Fig. 5, D-F), and CCL5-induced gene expression of *Vegf*, *Plat* and *Plau* was abrogated by the treatment of ERK inhibitor (Fig. 5, G-I). Collectively, these observations demonstrated that the CCL5-CCR5 axis could up-regulate VEGF, PLAT and PLAU by activating ERK pathway.

**Figure 5.**
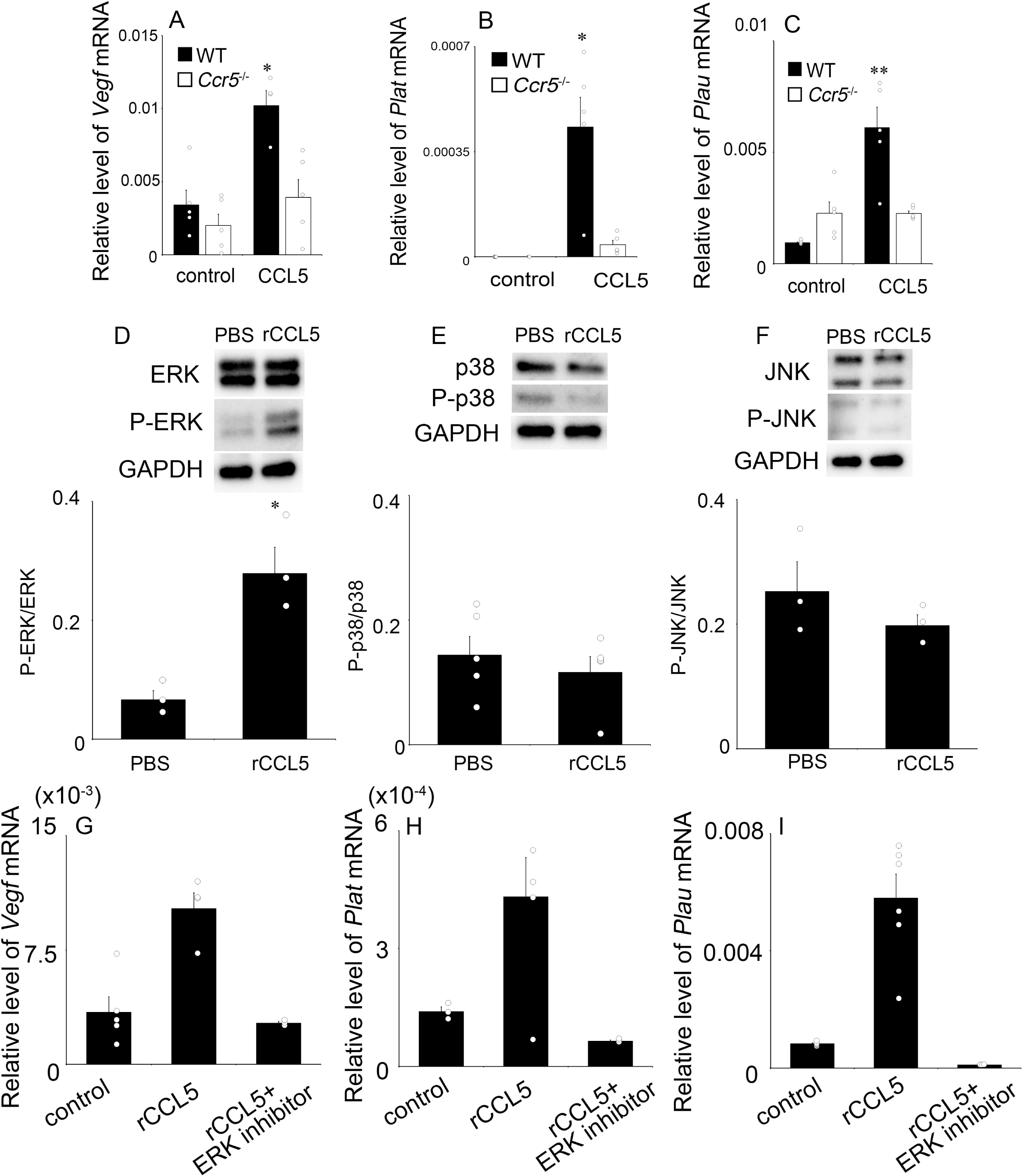
The effects of CCL5 on phosphorylation of ERK, p38 MAPK and JNK on peritoneal macrophages. (A-C) The effect of CCL5 on the gene expression of *Vegf* (A), *Plat* (B), and *Plau* (C) in peritoneal macrophages. Peritoneal macrophages were obtained from WT and *Ccr5*^-/-^ mice, and were stimulated with rCCL5 or PBS. All values represent the means ± SEM (5 independent experiments). \*\**P*<0.01; \**P*<0.05, vs. no stimulation. (D-F) Western blotting analysis using anti-GAPDH antibody confirmed that an equal amount of protein was loaded onto each lane. Representative results are shown. The ratio of phosphorylated (p)-ERK/ERK (D, 3 independent experiments), p-p38/p38 (E, 5 independent experiments) and p-JNK/JNK (F, 3 independent experiments) were densitometrically determined and are shown. All values represent means ± SEM. \*\**P*<0.01; \**P*<0.05, vs. no stimulation. (G-I) The effects of ERK inhibitor on CCL5-induced gene expression of *Vegf* (G, 4 independent experiments), *Plat* (H, 5 independent experiments) and *Plau* (I, 6 independent experiments). Each gene expression was analyzed by real-time RT-PCR. All values represent means ± SEM. \*\**P*<0.01; \**P*<0.05, vs. CCL5 stimulation alone.

## Discussion

Upon binding with a corresponding receptor, chemokines crucially regulate leukocyte migration and some of them additionally act to promote angiogenesis. Mirroring the fact that leukocyte infiltration and angiogenesis are major histological changes in venous thrombus formation and resolution, venous thrombus resolution is regulated by chemokines such as CXCL8 and CCL2, which can affect neutrophil and monocyte infiltration (17, 18, 36). CCR5 is expressed mainly by monocytes and macrophages as well as T lymphocytes and NK cells (37, 38), and has three specific ligands, CCL3, CCL4, and CCL5. IVC ligation induced CCR5-expressing cells and enhanced the expression of CCL5 in venous thrombi. Moreover, IVC treatment induced *Ccr5*^-/-^ mice to develop larger thrombus mass than WT mice. Moreover, the administration of a specific CCR5 inhibitor to WT mice recapitulated similar phenotypes as *Ccr5*^-/-^ mice while that of CCL5 caused the opposite phenotypes. Thus, the CCR5-CCL5 axis can attenuate thrombus formation. However, as there was no significant difference in the blood coagulability between WT and *Ccr5*^-/-^ mice, the absence of CCR5-mediated signals can have few, if any, effects on blood coagulability, the first step of thrombus formation.

Leukocyte recruitment is a hallmark of thrombus resolution as well as wound healing (39, 40). The magnitude of intrathrombotic leukocyte recruitment was proved to correlate with thrombus size (15). Moreover, anti-neutrophil serum administration induced neutropenia and eventually retarded venous thrombus resolution (41, 42). Furthermore, profibrolytic roles of macrophages were proposed on several observations (43–45). Actually, CD11b^+^Ly6C^Lo^ macrophages were proved to be crucial to venous thrombus resolution, due to their pro-reparative function (44). However, there are discrepancies on the roles of lymphocytes in thrombus resolution. The depletion of CD4^+^ and CD8^+^ T cells impaired thrombus resolution in mice (46), whereas NK cells and effector memory T cells inhibited thrombolysis by producing IFN-γ (13, 47, 48). CCR5 is presumed to regulate the recruitment of macrophages, T cells and other bone marrow-derived cells (11, 26, 49). Moreover, the analyses using bone marrow chimeric mice revealed that CCR5-expressing bone marrow-derived cells were crucial for thrombus resolution. Thus, we assumed that CCR5 deficiency retarded thrombus resolution by attenuating intrathrombotic recruitment of leukocytes including bone marrow-derived cells. However, there were no significant differences in intrathrombotic leukocyte recruitment between WT and *Ccr5*^-/-^ mice after IVC ligation. Thus, CCR5-mediated signals might contribute to thrombus resolution with few effects on leukocyte recruitment, which might be regulated by other chemokines than CCR5 ligands and/or other proinflammatory cytokines.

MMPs are a representative family of proteolytic enzyme and their enzymatic activities are proposed to be closely associated with thrombus resolution (13, 50). Although CCR5^+^ cells were identified as the main source of MMPs (28, 51), there is a discrepancy in the effects of CCR5-mdiated signals on MMPs expression. Blocking CCR5 down-regulated MMP-9 expression in rat glial cells (51), whereas CCL3 suppressed MMP-9 expression via CCR5 on macrophages in aortic aneurysm (28). We proved that the gene and protein expression of MMP-2 and MMP-9 were enhanced to similar extents in WT and *Ccr5*^-/-^ mice after IVC ligation. Thus, MMP-2 and MMP-9 have few roles in CCR5 deficiency-mediated impaired thrombus resolution.

Plasmin is generated from plasminogen by the action of tPA and uPA (52). This pathway is presumed mainly to regulate fibrin deposition in vascular trees and to activate pericellular fibrinolysis, which is required for cell migration in tissues. Singh and colleagues (53) demonstrated that the absence of uPA impaired thrombus resolution, indicating the essential involvement of plasmin system in thrombus resolution. We revealed that intrathrombotic levels of uPA and tPA, but not PAI-1, were significantly attenuated in *Ccr5*^-/-^ mice compared with WT mice. Moreover, uPA and tPA expression was detected in CCR5^+^ F4/80^+^ macrophages. Thus, CCR5 deficiency may deprive F4/80^+^ macrophages of the capacity to express uPA and tPA.

Neovascularization is additionally presumed to have a crucial role in thrombus resolution (17, 54). This notion is further strengthened by the observations that impaired intrathrombotic neovascularization delayed thrombus resolution (16, 32). Neovascularization is accelerated by several growth factors, particularly VEGF. Indeed, *Tlr4*^-/-^ mice exhibited impaired resolution of stasis-induced venous thrombosis along with decreased intrathrombotic VEGF expression (55). Moreover, adenovirus-mediated transfection of the VEGF gene enhanced thrombus recanalization and resolution in rats (56). Here, we demonstrated that CCR5 deficiency reduced VEGF levels in thrombi and attenuated neovascularization, thereby decelerating thrombus resolution. Furthermore, the main source of VEGF was identified as CCR5^+^ F4/80^+^ macrophages in thrombi. Thus, CCR5^-^ macrophages can infiltrate into thrombus but may be incapable of producing VEGF, similarly as observed on tPA and uPA expression.

Intrathrombotic CCL5 expression prompted us to evaluate the effects of CCL5 on the capacity of F4/80^+^ macrophages to express VEGF, tPA, and uPA. Consistently with the observation that the CCL5-CCR5 axis can activate human and mouse macrophages in an ERK-dependent manner (57, 58), CCL5 enhanced gene expression of VEGF, tPA, and uPA in WT-derived macrophages, but not *Ccr5*^-/-^-derived macrophages, by activating the ERK pathway. Thus, CCR5^-^ macrophages are not intrinsically defective in VEGF, uPA, and tPA expression but are unresponsive to CCL5 present in thrombus and as a consequence, failed to promote thrombus resolution.

In summary, CCR5-deficient mice exhibited reduced expression of *Vegf, Plat,* and *Plau,* with concomitant attenuated neovascularization and reduced thrombus resolution. the Thus, the augmentation of the CCL5-CCR5 axis may be effective for DVT treatment by enhancing thrombolysis. Moreover, as Ccr5Δ32 alle is a common loss of function allele in Caucasian population with a hemizygotic frequency of about 10 % (59–61), the individuals with this allele may be vulnerable to DVT and therefore, may have to be treated intensively in order to prevent serious sequelae such as PE and post-thrombotic syndrome.

## Data Availability

The authors declare that all data are available in the article file and Supplementary information files, or available from the authors upon reasonable request.

## Supplemental Material

Figures S1 and S2.

